# Mechanistic insights of the attenuation of quorum-sensing-dependent virulence factors of *Pseudomonas aeruginosa*: Molecular modeling of the interaction of taxifolin with transcriptional regulator LasR

**DOI:** 10.1101/500157

**Authors:** Hovakim Grabski, Susanna Tiratsuyan

## Abstract

Pseudomonas aeruginosa is one of the most dangerous superbugs and is responsible for both acute and chronic infection. Current therapies are not effective because of biofilms that increase antibiotic resistance. Bacterial virulence and biofilm formation are regulated through a system called quorum sensing, which includes transcriptional regulators LasR and RhIR. These regulators are activated by their own natural autoinducers. Targeting this system is a promising strategy to combat bacterial pathogenicity. Flavonoids are very well known for their antimicrobial activity and taxifolin is one of them. It is also known that flavonoids inhibit *Pseudomonas aeruginosa* biofilm formation, but the mechanism of action is unknown. In the present study, we tried to analyse the mode of interactions of LasR with taxifolin. We used a combination of molecular docking, molecular dynamics simulations and machine learning techniques, which includes principal component and cluster analysis to study the interaction of the LasR protein with taxifolin. We show that taxifolin has two binding modes. One binding mode is the interaction with ligand binding domain. The second mode is the interaction with the “bridge”, which is a cryptic binding site. It involves conserved amino acid interactions from multiple domains. Biochemical studies show hydroxyl group of ring A in flavonoids is necessary for inhibition. In our model the hydroxyl group ensures the formation of many hydrogen bonds during the second binding mode. Microsecond simulations also show the stability of the formed complex. This study may offer insights on how taxifolin inhibits LasR and the quorum sensing circuitry.

## Introduction

Antibiotic resistance crisis is worsening, so urgent action has to be taken to fix this problem. *Pseudomonas aeruginosa* is one of the ESKAPE pathogens and affects cystic fibrosis sufferers, burn victims and patients with implanted medical devices [1]. One potential strategy is to target the quorum sensing system, which is responsible for the cell-cell communication of the bacteria. This system regulates gene expression depending on the cell population density [1]. Bacteria and fungi produce signalling molecules, termed autoinducers (AI), which regulate the QS system[2]. Virulence factors, coordination between microorganisms and the microorganism and the host are regulated by autoinducers [2]. It has been shown that quorum sensing plays an essential role in *P.aeruginosa*’s virulence [2]. Quorum sensing also activates the CRISPR-Cas immunity system, thus suppressing it would make *P.aeruginosa* more susceptible to phage therapy and antibiotics [3]. Quorum sensing relies on proteins that synthesise and recognise AI. There have been determined four main QS systems so far, specifically the LasI/LasR and RhlI/RhlR [4], the PqsABCDE/PqsR [5] and AmbBCDE/IqsR [6]. Each QS system has its own AI’ and these different QS systems adjust and control each other [2]. So LasI/LasR regulate all the other three systems. Transcriptional regulator LasR is one of the main quorum sensing regulatory protein and it is activated by its autoinducer (AI), the N-3-oxododecanoyl homoserine lactone (3OC12-HSL) (Fig. 1).

**Fig 1.**
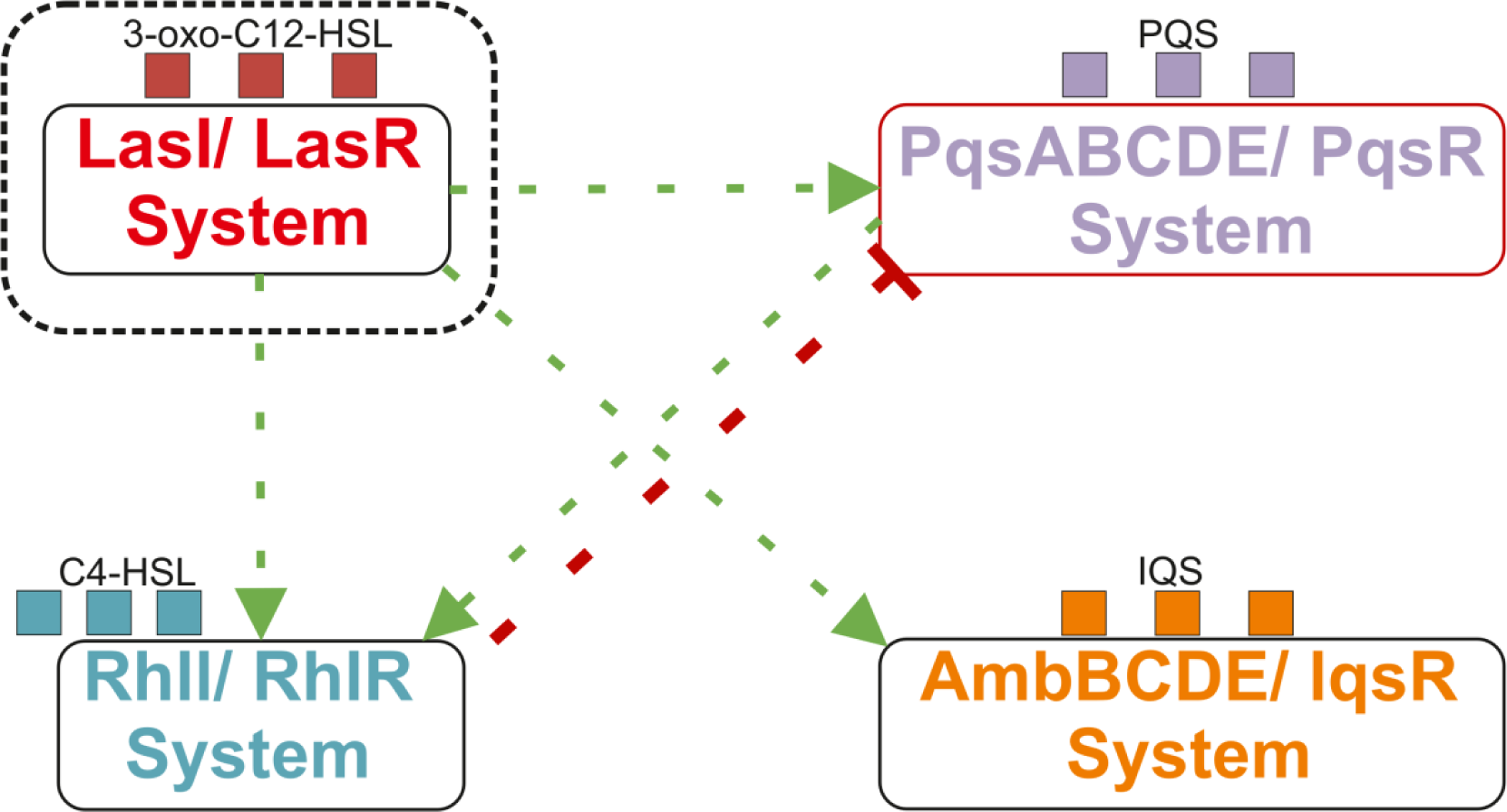
Quorum Sensing network of *P. aeruginosa*. LasR system is activated by the 3OC12-HSL which activates virulence gene expression and biofilm formation. Note: Green arrows depict activation and red arrows depict inhibition.

This protein is also activated by other homoserine lactones, which forms part of the interbacterial communication. LasR protein binds its AI, dimerizes, binds to promoter region of the DNA and activates gene expression. LasR protein has two domains: ligand binding domain (LBD) and DNA binding domain (DBD). One issue with this protein is that the whole structure is not available, because it is insoluble. Thus the inhibiting of LasR protein will prevent the activation of virulence genes and stop biofilm formation.

Flavonoids are a group of natural products and are known as Quorum quenching (QQ) agents [1]. It has been reported that flavonoids and their derivatives inhibit biofilm formation of *P.aeruginosa* [1]. Quorum quenching agents focus not on the killing of the bacteria, but reduction of virulence [2]. This approach does not cause antibiotic resistance, and it can be used in combination with other drugs. It has been shown that flavonoids inhibit LasR protein, but the mechanism of action is not known [1].Their results obtained through biochemical studies show that flavonoids inhibit through a non-competitive mechanism. They do not disrupt the formation of the dimer. They prevent LasR from binding to promoter region of the DNA. Flavonoids can bind to LasR LBD when AI is bound. They deduce that flavonoids do not use the canonical AI binding site. The authors also note that at low concentrations and only in the presence of the AIs, the flavonoids modestly activate the reporter strains. They also show that the OH group in position 7 of flavonoids is essential for inhibition [1]. In another work the authors [7] reveal that multiple antagonists bind to LasR, stabilize it and provoke an unnatural fold, which does not allow the binding of LasR to promoter sequence, which does not affect the dimerization as well. They also provide additional evidence that LasR-ligand complexes can be reversible. In our previous work we modelled the interaction of 3OC12-HSL with LasR [8]. It appeared there could be a possible second binding site. In another work we show that quercetin has two binding modes: one with LBD and the other with the “bridge” [9]. It has been shown that the LBD is capable of having multiple ligands in the binding pocket [10], but yet flavonoids do not inhibit in the presence of 3OC12-HSL [1].

It has been shown that the inhibitors do not affect dimerization interface [1, 7], so there has to be another mechanism for inhibition. Taxifolin is also a flavonoid, and it has been shown that it is not mutagenic less toxic than quercetin [11]. We show that taxifolin also has two binding modes, one with the LBD and the “bridge”, which includes amino acids from LBD and DBD, like quercetin [8, 9].

The interaction of inhibitors with DNA binding domain has been demonstrated for another transcriptional regulator in *P. aeruginosa*. Protein ExsA regulates type III secretion system (T3SS), which sabotages the host cell [12] and this protein also has HTH motif like the LasR protein. The ExsA is a transcriptional activator and belongs to AraC family, which regulates the operons encoding genes of the structural components of T3SS. Generally binding of arabinose to AraC triggers a rearrangement of an amino terminal extension, which in its turn permits the AraC dimer to bind to two adjacent binding sites on the promoter and activate transcription [13] (Fig. 1). In the case of ExsA it is regulated through protein-protein contacts rather than small molecules. The inhibitors of DBD of ExsA belong to N-Hydroxybenzimidazole [14]. As mentioned above the DBD of these two proteins contain HTH motif, so there could be a common mechanism. This could mean that there is a more nuanced modulation of the LasR and the ExsA.

As it has been shown that hydroxyl group at position 7 in flavonoids is essential for inhibition [1], we removed the hydroxyl group and ran another set of simulations. In this work we used a combination of molecular docking, molecular dynamics (an aggregate of 7.6 microseconds of simulations) and machine learning techniques, which include principal component and cluster analysis.

## Methods

### Entire flowchart

The whole methodology is presented as a flowchart for a better comprehension, with more details in the corresponding subsections:

- The structure of the LasR monomer is based on our previous research [8]. The model was validated using Gaia [15], QMEAN [16], PROSA [17], Verify3D [18].
- The 3D model of taxifolin was acquired from PubChem [19] web server.
- Ligand parameters generated using acpype [20] interface in the framework of the AMBER force field.
- Docking of taxifolin with the LasR monomer performed using Autodock Vina [21], LeDock [22], FlexAid [23], and rDock [24].
- Identification of binding sites from the docking results using Principal component and cluster analysis.
- Extraction of the centroid poses (the most likely conformation) from the clusters.
- Using the centroid pose as input for molecular dynamics simulations.
- MD simulations performed using Gromacs 2018 [25] with AMBER99SB-ILDN forcefield [26].
- PCA and cluster analysis of the MD trajectories.
- Binding energy calculation using MM-PBSA approach and sequence conservation analysis.
- Protein-protein docking using ClusPro [27].

### Validation of the protein model

The model was taken from our previous research [8] and was validated using GAIA [15], which compares intrinsic structural properties of the protein to experimental data (S1 Fig). It was also validated with QMEAN with a Z-score less than 1[16] (S2 Fig) and PROSA with a score of -7.17 [17]. Ramachandran plots showed no amino acids were in disallowed regions [8] and Verify3D indicated that 92.89% of the residues have averaged 3D-1D score >= 0.2 [8]. All these tools demonstrate that the model is of satisfactory quality.

### Ligand Molecule

The molecule (Fig. 2) was obtained from Pubchem [19]. The ligand was parameterised using the acpype [20] tool for the GAFF force field [28]. It has been shown that the OH group at position 7 is necessary for inhibition [1], so we created another ligand by removing this group using MarvinSketch [29] and replacing it with a hydrogen atom.

**Fig 2.**
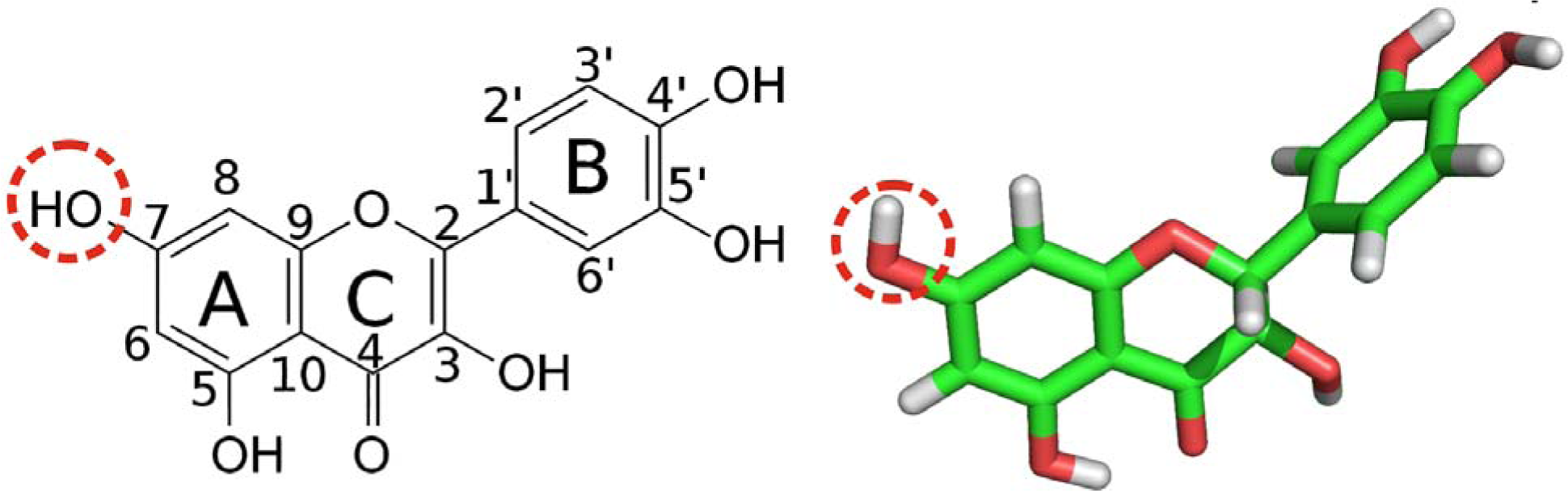
Taxifolin molecule structure. left) 2D right) 3D stick representation.

### Centroid (“Representative structure”) extraction

The centroid or representative structure was extracted using the following algorithm:

1. Compute all pairwise root mean square deviations (RMSDs) between the conformations.
2. Transform the distances into similarity scores and they are calculated as

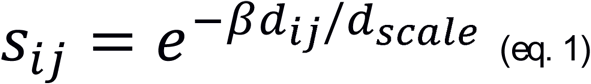

Where s_ij_ is the pairwise similarity, d_ij_ is the pairwise distance, and d_scale_ is the standard deviation of the values of d to make the computation scale invariant.
3. Then the centroid is defined with β=1 as

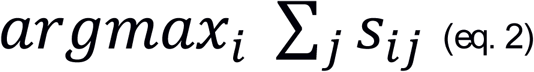

### Clusterization metrics

Clusterization is an unsupervised machine learning technique, but the determination of the number of clusters is not an easy task. For this purpose different metrics have been developed to solve this problem. Various metrics were used such as the David-Bouldin Index [30], Dunn Index [31], Silhouette score [32] and Calinski Harabasz [33], because they are widely used in the machine learning community. But each individual method has a drawback, because a good value does not imply best information retrieval. These metrics have their particular properties: the smaller DBI index is better and vice versa for the dunn, silhoutee and Calinski Harabasz give indication of the quality of clustering. The final number of cluster was selected using these 4 criteria by selecting the 3 best scores for each respective metric. For each place a score is assigned, the first place gets a score of 1, 2^nd^ - 0.85 and 3^rd^ – 0.5. In the end is selected the number of clusters with the highest score.

### Ligand docking using multiple docking programs

Blind docking was performed using multiple docking programs. Each program has its own bias, thus using multiple docking programs allow to compensate for their individual deficiencies. This approach imitates ensemble machine learning. Autodock Vina [21], rDock [22], and LeDock [23] were used based on their accuracy [35] and FlexAid [24] because of its stability against structural variance of the proteins [24]. A total of 40 docking trials were performed, 10 for each docking program. Each docking run was set to generate 20 poses. For Autodock Vina a grid box of 62*65*44 Å was chosen to cover the whole protein surface. The molecule for Autodock Vina was prepared using MGLTools. One run using the following exhaustiveness values: 8,16,32,64,128,256,512,1024,2048,4096 were performed for Autodock Vina. The radius was set to 36 Å for rDock and for LeDock the xmax was set to 103, xmin=41, ymax=105, ymin=40, zmax=56, zmin=12 to cover the whole protein surface. After the completion of docking simulations, the results were parsed with OpenBabel [35].

OpenBabel[35] is a chemical toolbox, which allows to parse many different types of chemical data. Later the coordinates of the center of mass of docking poses were subjugated to Principal component [36] and cluster analysis. They were performed using the scikit-learn library [37] for python programming language. The first two principal components were used for cluster analysis, which was performed using the DBSCAN algorithm [38]. This is a well-known algorithm and is used for data mining and machine learning [39]. It takes two parameters eps and minPoints. Eps was determined by plotting the kNN distance plot (S3, S4 Fig) Minimum number of points was set up to 5. After that only common clusters were selected, where all programs predicted a binding conformation. The centroid conformations were extracted from each cluster using equation 2:s

### Molecular dynamics simulations using the centroid docked poses

Gromacs 2018 [25] with AMBER 99SB-ILDN [26] force field were used for MD simulations. Steepest descent was used for minimization and was set Fmax of no greater than 1000 kJ mol-1 nm-1. Temperature was set to 300 K. In all experiments, the timestep was set to 2 fs. The leap-frog integrator [40] was used for all the simulations. LINCS [41] constraint was used for the bonds, which allows to increase the timestep to 2 fs. Explicit water solvation with TIP3P [42] waters was used. Van der Waals cut-off distance was set to 1.2 nm. Periodic boundary conditions were used for all the simulations.

In all simulations the concentration of NaCl was about 100 mM. In all cases, the short range non bonded interactions were truncated to 1.2 nm. Particle Mesh Ewald [43] was used for long range electrostatics. Ligand parameters generated using the acpype [20] tool. It generates GAFF compatible topology. After minimization, the system was equilibrated in 2 stages: the first stage involved simulating for 200 ps under a constant volume (NVT) ensemble, the second stage involved simulating for 200 ps under a constant-pressure (NPT) for maintaining pressure isotropically at 1.0 bar. Molecular dynamics trajectory was analysed with MDTraj [44] and scikit-learn [37]. Agglomerative clustering was used for the cluster analysis of MD trajectories, because this algorithm does not provoke a bias. To assess the quality of the clusterization, multiple metrics were used that are described in Methods section. To fit the trajectory in the memory for analysis a value of 12 was used for the stride. This means every snapshot apart 120 ps was used for the analysis. Hydrogen bonds were calculated using the Wernet-Nilson criteria [45], which can be calculated using the available module in MDTraj [44]. Final conformation was obtained by calculating the average structure using the last 500 frames. Binding energy was calculated using g_mmpbsa [46], using the last 20 ns from the trajectory. MMPBSA approach [47] was used because it provides good balance between accuracy and speed. It should be noted that MMPBSA approach can calculate the relative binding energy and not the absolute binding energy. For each interaction 167 snapshots were used for MMPBSA calculation.

### Protein-protein docking

When docking homology models, it is best if there is an experimental evidence to suggest the general interaction site (within ∼10 Å). Representative structures from molecular dynamics simulations were used for protein-protein docking using ClusPro [27]. ClusPro was chosen, because it was equivalent to the best human predictor group according to the latest CAPRI experiments carried out in 2013 [48]. From the experimental X-ray data, it was found that ‘Trp152’, ‘Lys155’ and ‘Asp156’ play an important role during dimerization. The distances between ‘Trp152’ from chain A and ‘Asp156’ from chain B was 0.276 nm, ‘Asp156’ from chain A and ‘Lys155’ from chain B of the crystallographic structure was 0.279nm. These residues were used as attraction constraints. The multimer docking mode of Cluspro was used for the prediction of dimer conformations.

## Results

### Docking of taxifolin with LasR

The methodology was inspired by our previous works [8, 9]. Molecular docking was performed using multiple docking programs: Autodock Vina [21], rDock [22], LeDock [23], FlexAid [24]. Multiple docking programs allow to obtain a consensus. For each program 10 simulation runs were performed. Later PCA analysis [36] was performed using scikit-learn [37] on the center of mass coordinates (COM) of the docking results. The first two principal components explain 86.17% variance in data. The identification of binding sites was performed in two stages. The first stage involved clusterization using the DBSCAN algorithm (Fig. 3b). The second stage involved selecting clusters where all 4 docking programs are featured for a consensus (Fig. 3c). It appears there are 2 potential binding sites for taxifolin with and without the hydroxyl group. For taxifolin these 2 clusters feature 80.4% of all the docking poses (Fig. 3). For the taxifolin without the hydroxyl group these 2 clusters amount 73.8% of all the docking poses. These two clusters correspond to ligand binding domain, which corresponds to the experimental binding site and the other is the “bridge” region [8, 9]. Then we extracted the “centroid” (representative structure) from each cluster of the docking simulations using equation 2 for molecular dynamics.

**Fig 3.**
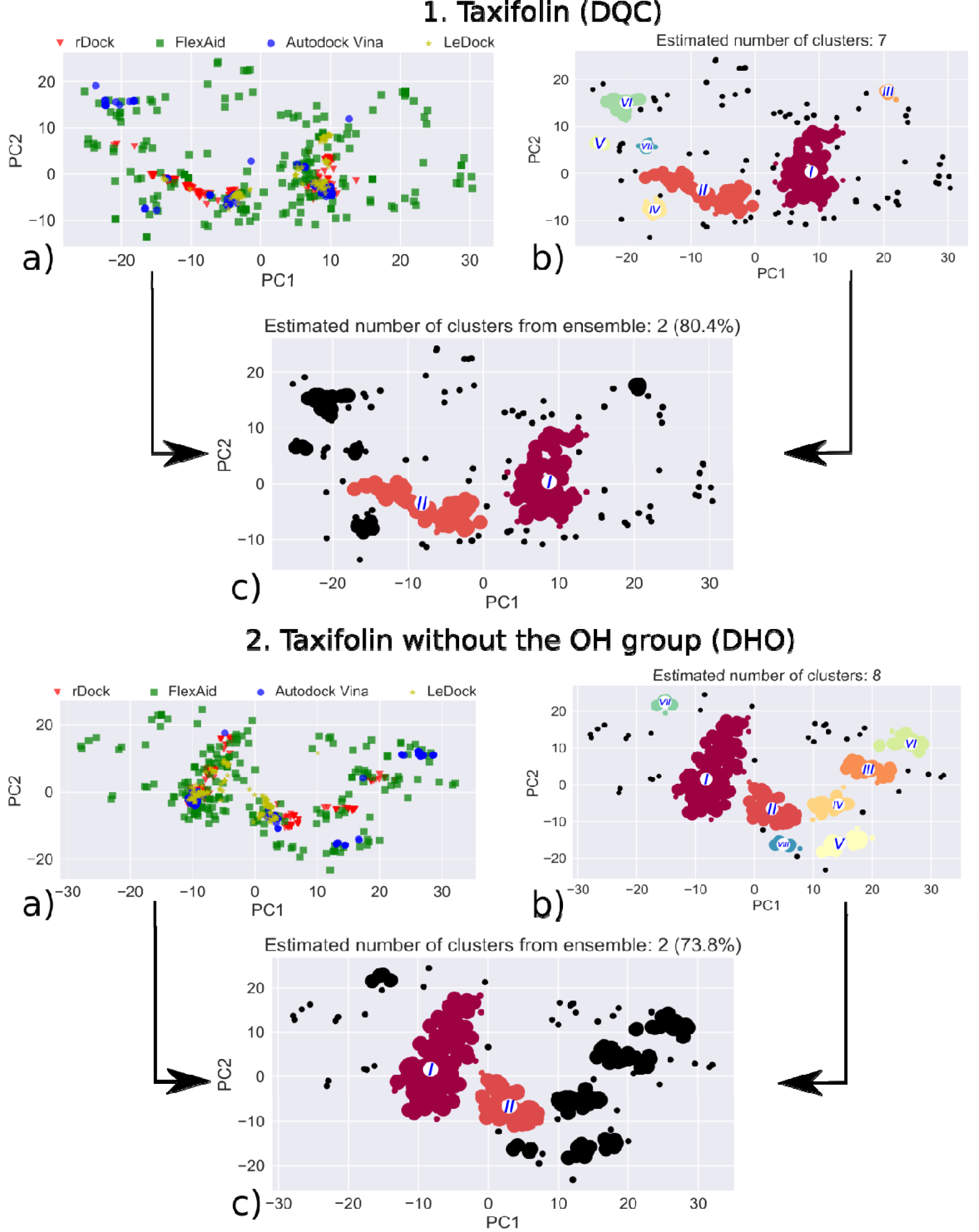
Analysis of blind docking of 1. taxifolin (DQC) and 2. taxifolin without OH group (DHO) with LasR. a) Visualization of the first two PCs of COM data from docking poses. b) Clusters c) Clusters, where all docking programs converged.

**Fig 4.**
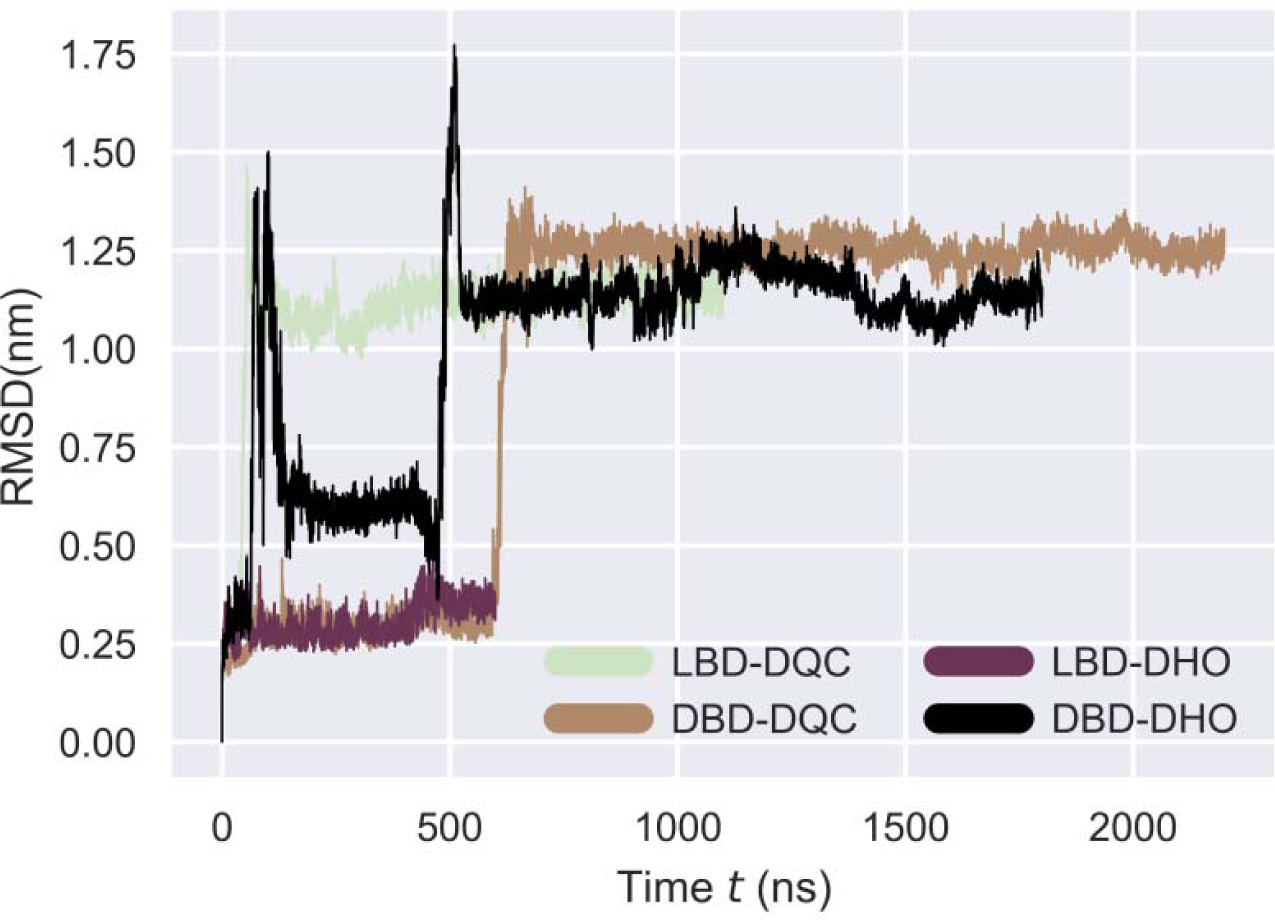
RMSD evolution through time for LasR monomer structure. The RMSD is calculated for the backbone atoms of the residues.

### Binding modes of taxifolin with LasR by MD simulations

The extracted centroid pose were used for molecular dynamics simulations. A total of 3.4 μs (DQC) simulations were performed for taxifolin (DQC). Taxifolin with the removed OH group (DHO), for each pose 5 simulations of 200 ns were run. From the 5 simulations for both cases, in one of the simulation the molecule continued interacting with LasR. These cases were extended to 1.8 μs for the “bridge” and 600ns for LBD. Overall, 3.4 μs (DQC) and 4.2 μs (DHO) of simulation data wa collected. The simulations were performed using Gromacs 2018 [25] with Amber ff99SB-ILDN [26] force field. The obtained data were used for the analysis of the interaction of DQC and DHO with LasR monomer. An aggregate of 7.6 microseconds of simulation data was used for the analysis of the interactions.

The structure of the LasR protein in all of the interactions appears to be stable and converged based on the RMSD plot (Fig. 4).

### Binding modes in the LBD

Taxifolin (DQC) does not enter the ligand binding pocket of the LasR like the native autoinducer 3OC12-HSL [8]. This interaction on the outer surface of the LBD provokes an allosteric change of the DBD. It forms only one hydrogen bond with Gln94 and interacts hydrophobically with conserved Glu103 and other residues (Fig. 5a). The DHO does not enter the binding pocket either but remains close to it (Fig. 5b).. It also forms one hydrogen bond but with Lys42 and does not interact with many conserved residues like DQC. It does not affect the conformation dramatically like DQC does.

**Fig 5.**
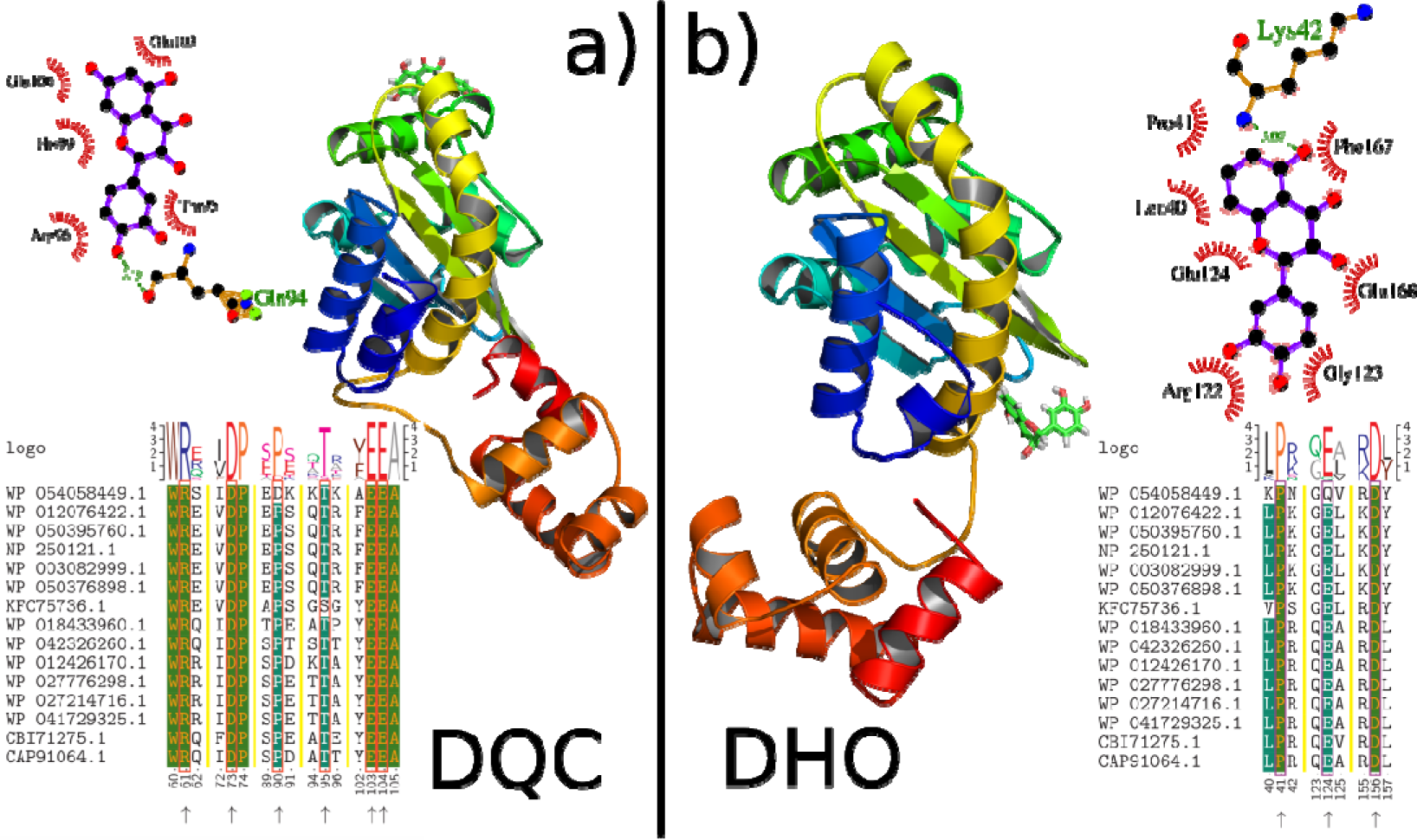
Binding modes of taxifolin (DQC) and taxifolin without OH (DHO) group to the LBD of transciptional regulator LasR. It also presents the respective hydrogen bonds and interactions with the conserved residues.

The relative binding energy shows that DQC has stronger relative binding affinity than DHO. The removal of OH group increased the van der Waals components, but reduced the electrostatic and polar solvation contribution (Table 1). Overall it appears the binding affinity of DHO is lower than DQC.

**Table 1.** Relative binding energy calculations using MMPBSA approach for LBD interaction using g_mmpbsa (kJ/mol).

| Binding sites | Van der waals | Electrostatic | Polar salvation | Non-polar | Binding Energy |
| --- | --- | --- | --- | --- | --- |
| HSL-LBD | -215.67±8.007 | -39.59±18.82 | 147.52±13.99 | -20.84±0.79 | -128.58±18.76 |
| DQC-LBD | -90.95±12.55 | -163.92±39.43 | 148.55±24.70 | -10.17±0.64 | -116.51±18.92 |
| DHO-LBD | -130.62±9.401 | -51.66±33.186 | 100.64±18.34 | -13.26±0.64 | -94.90±17.32 |

**Table 2.**
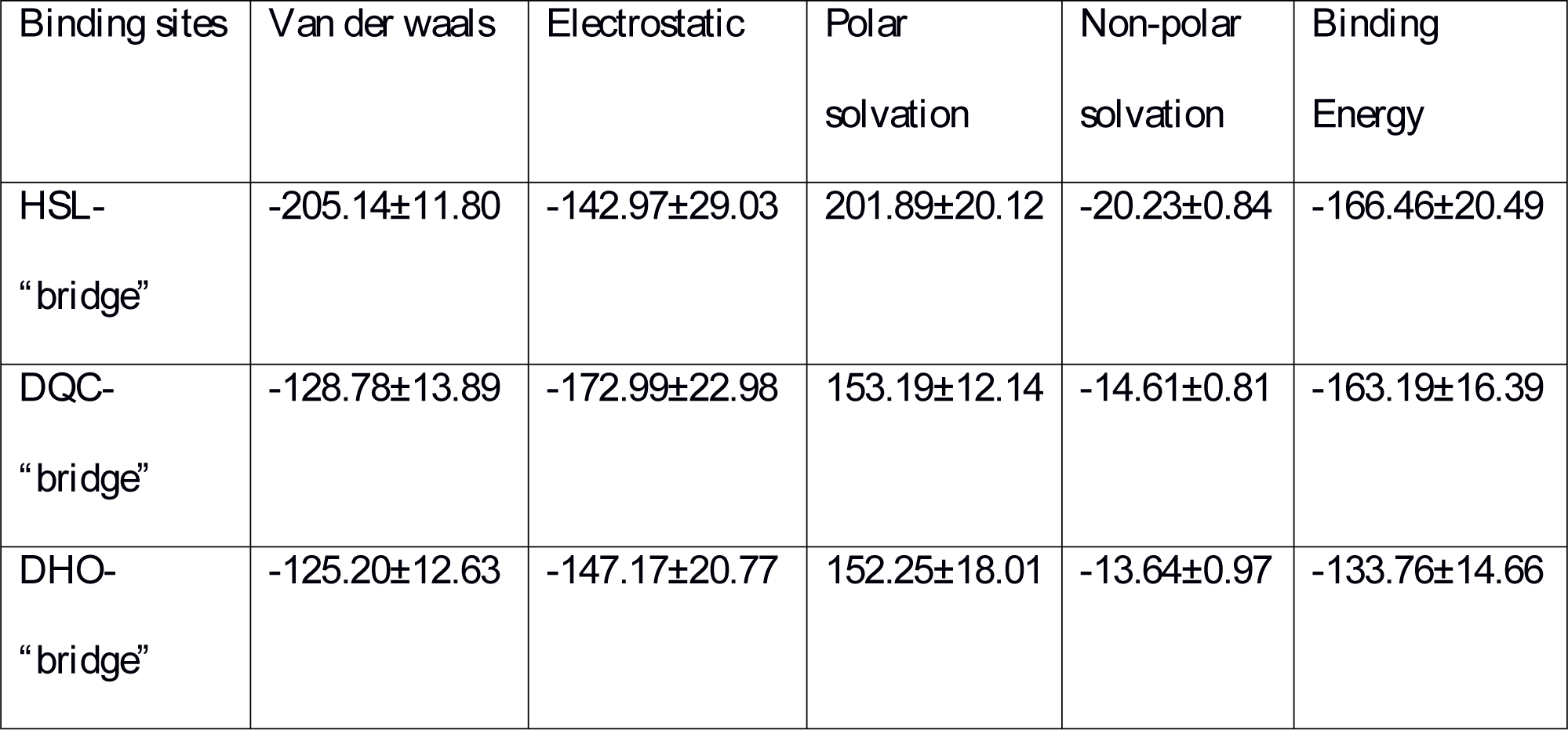
Relative binding energy calculations using MMPBSA approach for the “bridge” interaction using g_mmpbsa (kJ/mol).

### Binding modes in the the “bridge” DBD

Taxifolin (DQC) provokes major changes of the DBD, while DHO does not. The DQC forms 3 hydrogen bonds and interacts hydrophobically with Leu234, Glu168, Gln160, His169 and Val176 (Fig. 6a), while DHO forms 2 hydrogen bonds with Lys42 and Leu39 and interacts hydrophobically with Leu40, Pro41, Val171, Arg122 and Glu 124 (Fig. 6b). It should be noted that DQC interacts with many conserved amino acid residues, including Leu177 and Leu236. The native autoinducer [8] and quercetin [9] also interacts with Leu177 and Leu236, while DHO does not.

**Fig 6.**
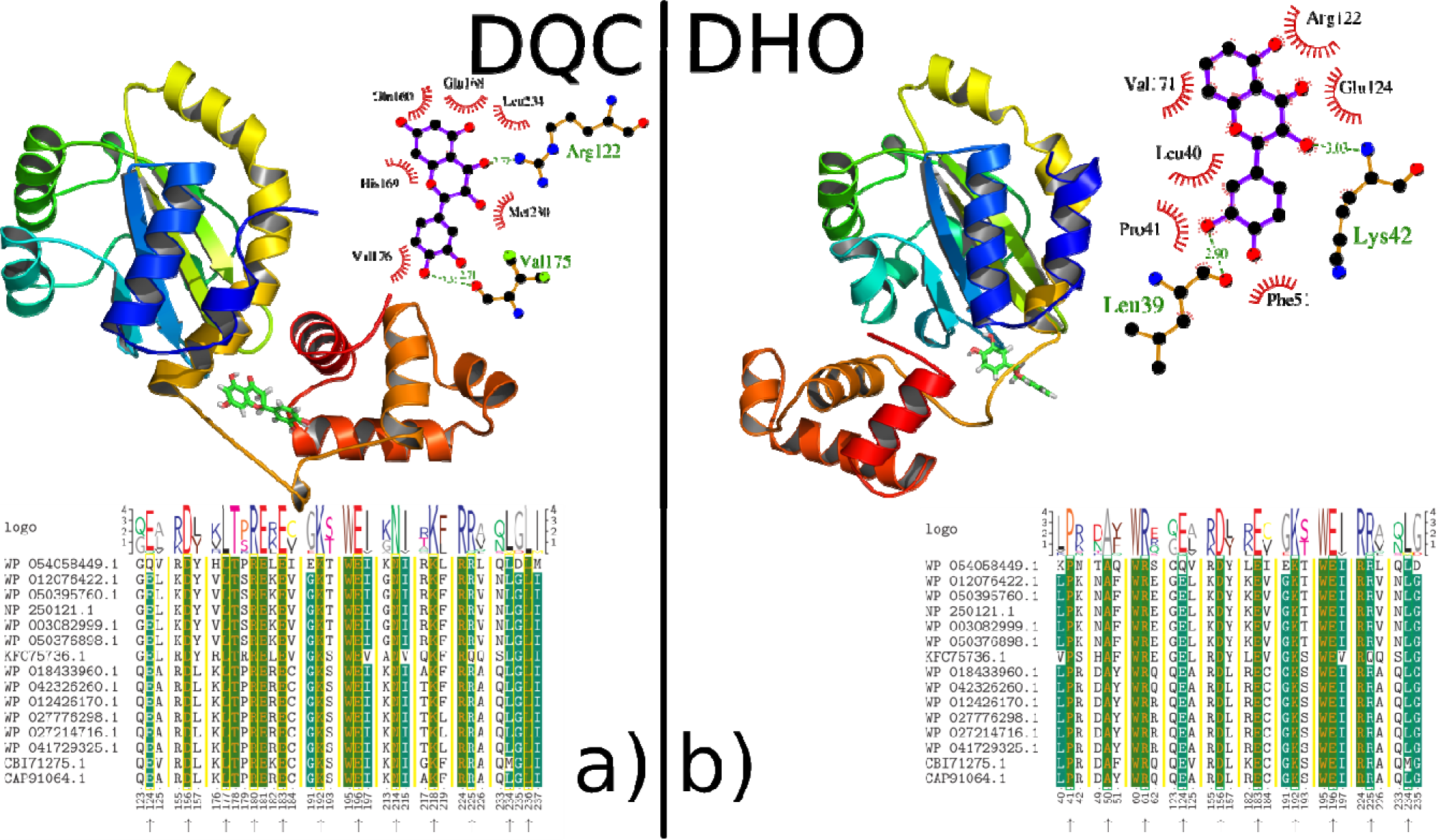
Binding modes of taxifolin (DQC) and taxifolin without OH (DHO) group to the LBD-SLR-DBD “bridge” (cryptic binding site) of transcriptional regulator LasR. It also presents the respective hydrogen bonds and interactions with the conserved residues.

The relative binding energy shows that DQC has stronger binding affinity than DHO. The removal of OH group decreased the electrostatic contribution (Table 1). Overall it appears the binding affinity of DHO is lower than DQC. The binding affinity of DQC is similar to the native autoinducer.

### Protein-protein docking

It has been shown that OH group at position 7 is essential for inhibition. Flavonoids do not inhibit dimerization, which potentially means that flavonoids do not interact with the dimerization interface [1]. It has also been suggested that inhibition happens through a non-canonical site [1]. Inhibition by the flavonoids is not competitive [1] and in the presence of 3OC12-HSL, they do not inhibit gene expression. It has also been shown that the inhibitors of LasR provoke an unnatural fold, yet do not affect dimerization [7] For that purpose we performed protein-protein docking using the conformations induced by DQC and DHO. The conformations are the average structure from the last 500 frames of MD simulations. ClusPro was used for protein-protein docking, since it has also been validated in the CAPRI challenge [27]. The closest homologue where the full structure is available to LasR, is the quorum sensing control repressor, QscR, Bound to N-3-oxo-dodecanoyl-L-Homoserine Lactone (Figure 7). The orientation of DBD are on the same plane. The docking conformations of the LasR-DHO, the orientation of DBD are also on the same plane thus can potentially interact with DNA. The docking conformations of the LasR-DQC complexes, the orientation of DBD are not on the same plane, thus this may explain how DQC stops the binding of LasR to promoter sequence.

**Fig 7.**
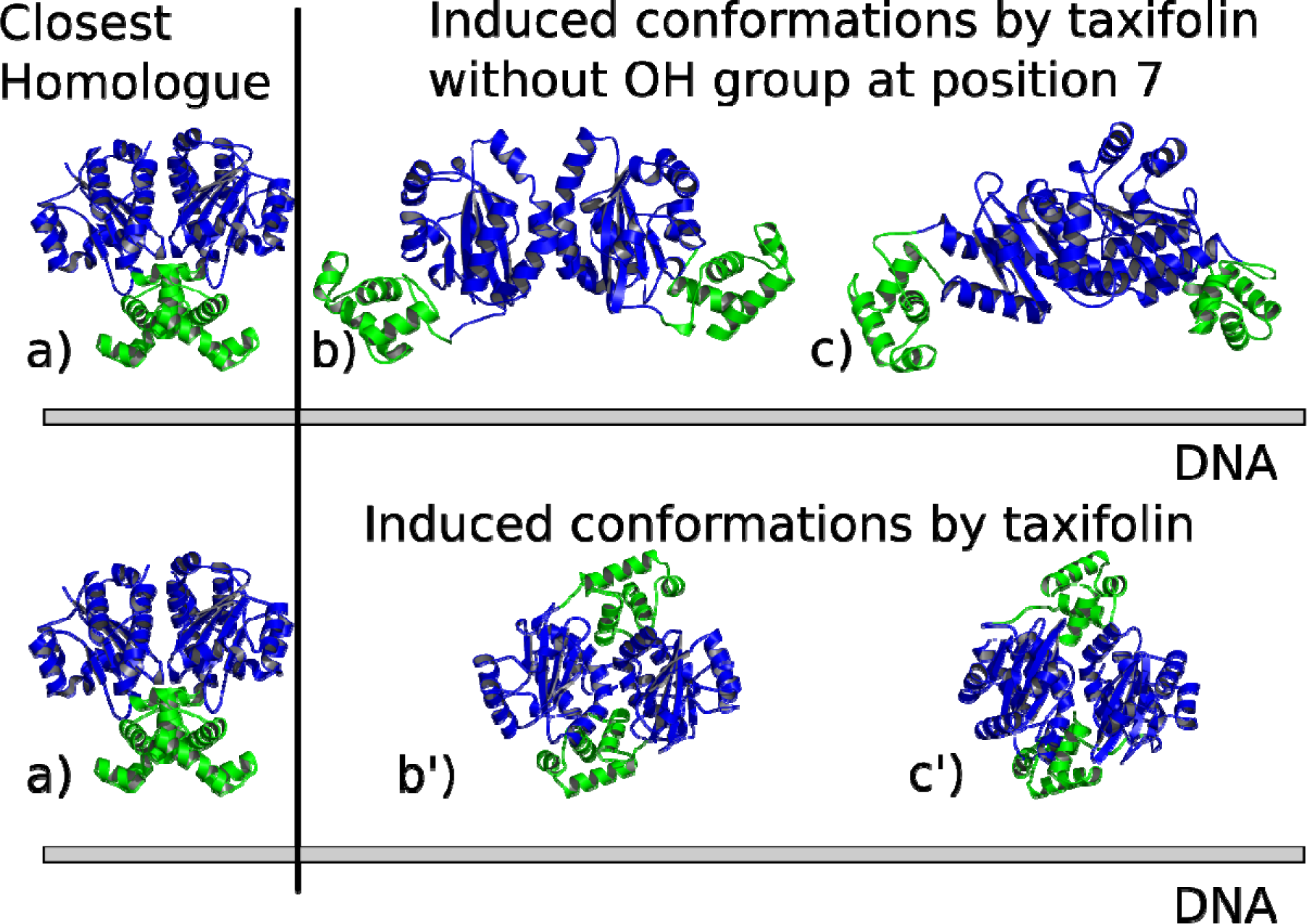
Comparisons of dimers: a) Quorum Sensing Control Repressor, QscR, Bound to N-3-oxo-dodecanoyl-L-Homoserine Lactone b) Induced conformations of the dimer by the binding mode of DHO with LBD c) Induced conformations of the dimer by the binding mode of DHO with LBD-SLR-DBD “bridge” b’) binding mode of DQC with LBD c’) Induced conformations of the dimer by the binding mode of DQC with LBD-SLR-DBD “bridge”.

## Discussion

*Pseudomonas aeruginosa* is a difficult bacterium to deal with and antibiotics are not effective against them. People are becoming more vulnerable to these bacteria. One potential way to counter is targeting the quorum sensing system. LasR protein has been known for a decade, albeit partially, because the full protein is not soluble. But yet, drug design has not been that prominent. Biochemical studies indicate the flavonoids do not affect the dimerization [1]. They do not interact directly with the DNA. It does not happen through the canonical site of the native autoinducer. The hydroxyl group at position 7 is essential for inhibition [1]. So for that purpose we emulated the biochemical study by removing the OH group from taxifolin. The induced conformation by taxifolin and subsequent protein-protein docking indicate that the DBDs do not face the same orientation as the DNA.

This may explain how flavonoids inhibit the LasR protein. As for the induced conformations of the LasR by taxifolin without the OH group, which does not inhibit, the DBD face to the same direction and plane as the promoter DNA. The direct interaction of DQC with the “bridge” of LasR is more suitable to the experimental data, because it appears that 3OC12-HSL can also interact with this region [7], so it is more stable. It should be noted that the interaction of inhibitors with the DNA binding domain is nothing new. Another transcriptional regulator of *P. aeruginosa*, the ExsA protein in particular, consists of LBD and DBD, which also has the HTH motif [11s]. ExsA is responsible for the activation of type III secretion genes, which sabotages the host cells. It has been shown that N-Hydroxybenzimidazoles inhibit the activation of T3SS via the inhibition of the DBD of this protein [13]. This could suggest that there could be an underlying common mechanism for the transcriptional regulators of *P. aeruginosa*.

## Conclusion

Biofilms are huge problem nowadays, starting from tooth plaques up to food processing and health issues. In *P. aeruginosa* the LasR protein is the main regulator of biofilm formation. Taxifolin binds to the LBD-DBD “bridge” of transcriptional regulator LasR and interacts with conserved amino acid residues. They include Leu177 and Leu236 like the native AI. The binding energies of the 3OC12-HSL and taxifolin to the LBD-DBD “bridge” of LasR are close. The removal of the OH group at position 7 in taxifolin reduces the binding affinity. Biochemical studies have also shown that this OH group is essential for the inhibitory activity This protein has been known for a decade but yet drug design targeting this protein has not been prominent. Considering it has been shown that it stops biofilm formation and makes bacteria vulnerable to antibiotics. Imitation of the experiment by removing the OH groups shows that the computational model does not counterdict it, but can give insight of the mechanism of inhibition.

The protein-protein docking experiment shows that removing the OH group does not affect the orientation of the DBD and thus can bind to DNA. This is compatible with the experimental data, where it has been shown that the OH group is essential for inhibiton. The addition of the OH group in taxifolin radically affects the conformation of the protein, thus it cannot interact with the DNA. Longer simulations demonstrate that the structures are stable. The OH group ensures the interaction with conserved amino acid residues like Leu177 and Leu236. It appears these two residues are essential for inhibitory activity. The allosteric interaction which leads to the change of the DBD conformation does not explain fully the experimental data. The biochemical study shows that in the presence of 3OC12-HSL, flavonoids do not inhibit, but can even activate it. The direct interaction of taxifolin with the “bridge”, which is not competitive as well, matches the experiments a lot better and fully. All these interactions show that they are not competitive and this matches the biochemical studies. This research could give a potential answer on how flavonoids inhibit biofilm, virulence gene expression, CRISPR-Cas adaptive immune system of *P. aeruginosa*.

## Supporting information

**S1 Fig. Summary of scores for the input structure from Gaia.**

**S2 Fig. Normalized QMEAN6 score of the input structure.**

**S3 Fig. kNN distance plot of the docking results of taxifolin (DQC) with LasR for the estimation of the eps parameter.**

**S4 Fig. kNN distance plot of the docking results of taxifolin without OH group (DHO) with LasR for the estimation of the eps parameter.**

## Declarations

### Acknowledgments

- This work was made possible by a research grant from the Armenian National Science and Education Fund (ANSEF) based in New York, USA.
- We gratefully acknowledge the support of NVIDIA Corporation with the donation of the Titan Xp GPU used for this research.
- The research was done within the Ministry of Education and Science of the Republic of Armenia, State Committee of Science: 10-2-1-4, Government budget financing.

